# Mapping of critical prosodic and phonetic networks in post-stroke apraxia of speech

**DOI:** 10.64898/2025.12.16.694760

**Authors:** G. Lynn Kurteff, Grant M. Walker, Julius Fridriksson, Gregory Hickok

## Abstract

**Purpose:** Many have made proposals to better diagnose and/or classify post-stroke apraxia of speech (AOS), with some arguing for the separation of AOS into behavioral subtypes. Recent studies of primary progressive AOS have promoted a separation of prosodic and phonetic subtypes, aligning with a dual-motor coordination model separating the neural substrates of prosodic and phonetic function. Motivated by the limited corroboration of these subtypes in post-stroke AOS, here we present mapping results in a cohort of stroke survivors aiming to identify distinct neural substrates for prosodic and phonetic aspects of speech motor coordination.

**Methods:** Left-hemisphere stroke survivors (*n* = 127; 64 with AOS) received speech-language evaluation and neuroimaging at the Center for the Study and Treatment of Aphasia Recovery (C-STAR). AOS severity was quantified via the Apraxia of Speech Rating Scale (ASRS). We utilized a novel lesion-symptom mapping technique with an emphasis on prediction that identifies ensembles of regions supporting performance in the prosodic and phonetic domains.

**Results:** An ensemble of networks supporting prosodic function localized to dorsal and ventral (but primarily dorsal) sensorimotor cortex, as well as a distributed network of white matter pathways connecting Rolandic cortex to auditory regions and cerebellum, emphasizing the role of auditory feedback processing and laryngeal control in supporting prosodic function. A separate but partially overlapping network supporting phonetic function localized primarily to ventral Rolandic cortex and the arcuate fasciculus.

**Conclusions:** This work represents the first mapping of prosodic and phonetic subtypes in post-stroke AOS in a large cohort of individuals. We hope our results motivate the development of assessment and treatment techniques individually targeting prosodic and phonetic functioning to better serve individuals with AOS and facilitate clinical discussion of the disorder.

## Introduction

Speech production is a complex, multistage process. Psycholinguists have identified at least three stages involving the access of conceptual semantic, word (or morpheme), and phonological information (Levelt, 1993). Beyond these stages, research on speech motor control has identified two additional broad stages, a premotor level involved in some form of speech planning or multi-effector coordination, and a lower, primary motor level involved in execution (Guenther, 2016; Hickok, 2012). A recent theoretical synthesis has argued that the post-linguistic levels of processing are subdivided into two parallel hierarchies, one for phonetic articulation and one for voice pitch and prosody control (Hickok et al., 2023). The phonetic articulation system is hypothesized to involve premotor cortex on the precentral gyrus (the ventral precentral speech area, vPCSA) coordinating orofacial motor cortex, while the pitch/prosody system is hypothesized to involve a more dorsal precentral speech area (dPCSA) located just posterior to the middle frontal gyrus and overlapping area 55b (Glasser et al., 2016).

As lesion studies remain one of the only causal methods in cognitive neuroscience, cohorts with impaired motor speech coordination following neurological impairment offer an opportunity to test this hypothesis of a dual motor coordination hierarchy. In particular, people with apraxia of speech (AOS), a communication disorder primarily characterized as a difficulty with motor coordination (Darley, 1968; Johns & Darley, 1970; McNeil & Kimelman, 2001), may show different profiles of behavioral impairment with corresponding damage to the different pathways of motor speech coordination proposed in Hickok et al., 2023. A few published case studies have shown that surgical resection in the dorsal precentral gyrus, near the dPCSA, results in apraxia of speech with noticeable prosodic deficits (Chang et al., 2020; p.c. with authors of Levy et al., 2023, December 16, 2024). Studies of prosodic and phonetic ability in people with post-stroke AOS are limited to a single study with a small sample size (*N* = 8) that did not divide the precentral gyrus into the proposed dorsal and ventral components representing the prosodic and phonetic elements of motor coordination (respectively), instead subdividing these functions along an anterior/posterior split in precentral gyrus (Takakura et al., 2019).

AOS is considered difficult to differentially diagnose from dysarthria and expressive aphasia (Kobayashi & Ugawa, 2013; Patidar et al., 2013; Polanowska & Pietrzyk-Krawczyk, 2016; Ziegler et al., 2012), in part due to disagreement surrounding the primary diagnostic criteria of AOS (Haley et al., 2021) and whether or not AOS is divisible into behavioral subtypes (Mailend & Maas, 2020). Of particular interest to the current study is the debate surrounding subtypes of AOS: if there are indeed distinct, separable subtypes of behavioral impairment present in what is currently referred to monolithically as AOS, such subtypes should have separable neurobiological patterns of impairment.

Two subtypes proposed in neurodegenerative primary progressive apraxia of speech (PPAOS) appear to align neuroanatomically and behaviorally with the prosodic and phonetic motor coordination hierarchy (Josephs et al., 2013; Utianski et al., 2018). *Prosodic* PPAOS is marked behaviorally by difficulties with speech rate and syllable segmentation and was neuroanatomically localized in this study to the dorsal supplementary motor area (SMA) and superior cerebellar peduncle. The focal area of atrophic overlap identified in Utianski et al., 2018 overlaps with the dPCSA presented in Hickok et al., 2023. *Phonetic* PPAOS, on the other hand, is marked behaviorally by distorted sound substitutions and a more distributed network of impairment to the lateral SMA and precentral gyrus. While Utianski et al., 2018 localized phonetic PPAOS to a more distributed network in sensorimotor cortex and the cerebellum, the ventral precentral speech area (vPCSA) is still implicated within the region of atrophic overlap.

With a growing body of evidence in the neurodegenerative literature for prosodic and phonetic subtypes of PPAOS, a recent theoretical model of motor speech aligning with this dichotomy, and small-sample/case study results from lesion studies, the stage is set for a more detailed analysis of prosodic and phonetic ability in people with post-stroke AOS.

The current study aims to provide just that: an exploration of the neural substrates of prosodic and phonetic AOS in a large cohort of stroke survivors. In doing so, we aim to provide evidence for the hypothesis of prosodic and phonetic motor coordination hierarchies and motivate the consideration of prosodic and phonetic subtypes in post-stroke AOS. Our cohort, collected through the University of South Carolina’s Center for the Study of Aphasia Recovery (C-STAR), consists of structural imaging and comprehensive behavioral assessment in 107 individuals with post-stroke aphasia as well as 20 control stroke survivors without communication disorders. The Apraxia of Speech Rating Scale (ASRS; Strand et al.,(Strand et al., 2014) served as the primary assessment of apraxia of speech in this dataset. We employed a novel lesion-symptom mapping technique with an emphasis on predicting behavioral scores in held-out data to explicitly test the hypothesis that prosodic ability should localize to the dorsal precentral gyrus near area 55b while phonetic ability should localize to the ventral precentral gyrus just posterior of Brodmann area 44.

## Methods

### Participants

Data in this study are pulled from C-STAR’s Predicting Outcomes of Language Rehabilitation (POLAR) study; see (Kristinsson et al., 2023) for a thorough discussion of the dataset. Specifically, this study makes use of the baseline assessment data, which were collected and administered by ASHA-certified speech-language pathologists. Participants with left hemisphere stroke and aphasia were recruited for the study along with 20 control participants who were stroke survivors but without aphasia (*N*=127; Table 1). Participants underwent comprehensive speech and language evaluation; the ASRS (Strand et al., 2014) served as the primary diagnostic measure for AOS; therefore, participants with incomplete ASRS evaluations (*N*=4) were excluded from this study. 61 of the non-control participants (57%) had apraxia of speech, a relatively high concentration; however, none of the participants had isolated AOS. This is not uncommon, as “pure” AOS is rare and mostly restricted in clinical discussion to case studies (Chang et al., 2020; Levy et al., 2023).

**Table 1.** Participant demographics.

| Characteristic (Categorical) | Category | Qty. (N = 123) | Percentage |
| --- | --- | --- | --- |
| Sex | Female | 56 | 45.5% |
|  | Male | 67 | 54.5% |
| Race | White | 90 | 73.2% |
|  | Black or African American | 32 | 26.0% |
|  | Asian | 1 | 0.8% |
| Stroke type | Ischemic | 76 | 61.8% |
|  | Hemorrhagic | 33 | 26.8% |
|  | Other | 14 | 11.4% |
| Gross pathology | Control (WNL) | 20 | 16.3% |
|  | Aphasia only | 35 | 28.4% |
|  | Aphasia & dysarthria | 7 | 5.7% |
|  | Aphasia & AOS | 39 | 31.7% |
|  | Aphasia, dysarthria, & AOS | 22 | 17.9% |
| Aphasia type (WAB) | Anomic | 29 | 23.6% |
|  | Broca's | 48 | 39.0% |
|  | Wernicke's | 5 | 4.1% |
|  | Conduction | 15 | 12.2% |
|  | Global | 5 | 4.1% |
|  | Transcortical motor | 1 | 0.8% |
| Characteristic (Numeric) | Mean ± SD | Range | |
| Age (at stroke onset) | 55.7 ± 11.8 | 27 - 79 | |
| Age (at assessment) | 60.4 ± 10.8 | 29 - 80 | |
| Months post stroke onset | 54.9 ± 54.8 | 10 - 241 | |
| Education (years) | 15.5 ± 2.4 | 12 - 20 | |
| WAB Aphasia Quotient | 65.7 ± 25.2 | 14.5 - 100 | |

### Behavioral Assessment

The original version of the Apraxia of Speech Rating Scale (ASRS-1; Strand et al., 2014) was administered to evaluate the presence or absence of apraxia of speech (Figure 1A). The ASRS does not have dedicated stimuli and is instead scored by the speech-language pathologist using all speech produced during other assessments and informal conversation. Items are scored on a five-point scale (0 to 4), with a higher value indicating more severe impairment (0 being the absence of impairment) in that domain. The scale is split into four sections: (1) primary distinguishing features of AOS; (2) distinguishing features unless dysarthria is present; (3) distinguishing features unless aphasia is present; and (4) distinguishing features unless dysarthria and/or aphasia are present, reflecting the historic “process of elimination” diagnostic method for AOS. These data were collected before a notable revision to the ASRS in 2023 (ASRS-3.5; Duffy et al., 2023), which contains essentially the same content (save for three removed items) but reorganized into new sections supporting the prosodic and phonetic subtypes of PPAOS: (1) phonetic features; (2) prosodic features; and (3) other (most in this category concern diadochokinetic rate, a common bedside evaluation for AOS). Because this updated organization is aligned with our research question, we opted to generate scores for the ASRS-3.5 categories using our ASRS-1 data (Figure 1B). This gives us subscores that correspond to prosodic and phonetic ability directly; our sample contains a mixture of participants with impairments in one, both, or neither of these domains (Figure 1D).

**Figure 1.**
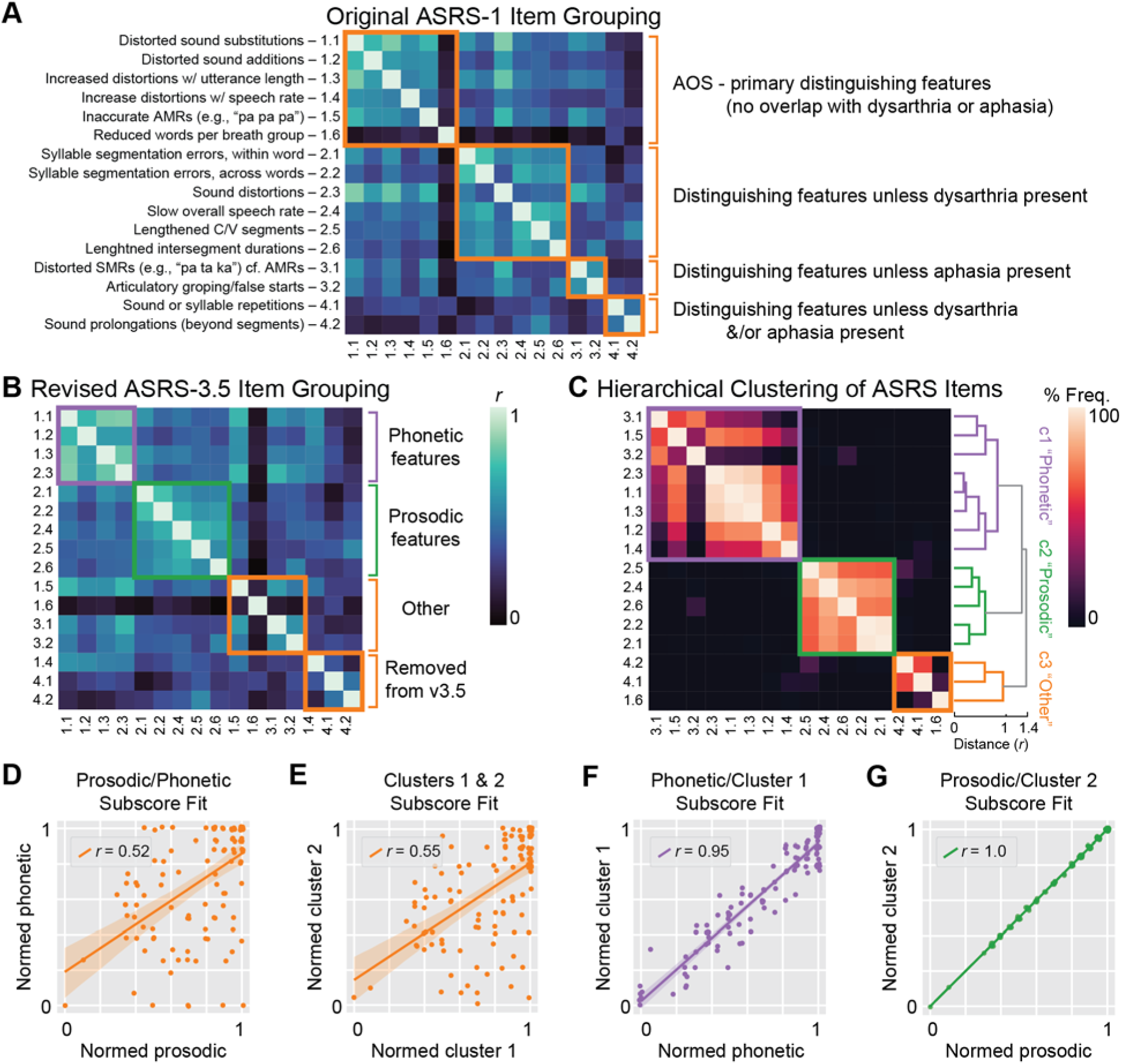
Constructing prosodic and phonetic subscores from Apraxia of Speech Rating Scale items. A: Confusion matrix of itemwise correlations on the ASRS-1. The four subsections of the ASRS-1 are emphasized using brackets to the right of the matrix. The original subgroups of the ASRS-1 do not always result in strong intra-group correlations across items; for example, item 1.6 does not correlate strongly with other items in “Primary features of AOS,” while 2.3 does correlate strongly with the items of this group but is in a separate group (“Features of AOS unless dysarthria present”). B: Another confusion matrix of itemwise correlations, but items are arranged based on subscores derived from ASRS-3.5. There are stronger intra-group correlations across items compared to the ASRS-1 subscores, but there are still some items that correlate strongly outside their group identity (e.g., items 1.5 and 3.1 from the “Other” group correlate moderately with items from the “Phonetic features” group). C: An alternative approach to ASRS subscore construction based on hierarchical clustering. The confusion matrix (left) shows frequency of co-occurrence of individual ASRS items in the same branch of an agglomeratively clustered dendrogram (right) using a bootstrapped distribution. The names assigned to the three clusters of the dendrogram are based on their resemblance to ASRS-3.5 subscores and reflect our hypotheses as to what these clusters are assessing. D: Regression plot showing the relationship between prosodic and phonetic subscores derived from the ASRS-3.5 grouping. Scores normalized and inverted so that a score of 0 indicates severe impairment while a score of 1 indicates little to no impairment. A small jitter (0.01) was introduced to point locations to better visualize individual subjects in categorical data. Single subjects (points) above the regression line have more relative impairment in phonetic function while subjects below the regression line have more relative impairment in prosodic function. There is a relationship between prosodic and phonetic subscores (normalized here to account for different maximum raw scores; Pearson *r* = 0.523; *p* < .001). This is not unexpected, as severe impairment in one domain progresses towards mutism, which impairs performance on all ASRS items to an extent. E. Regression plot similar to D showing the relationship between clusters 1 & 2 from the unsupervised method. F. Regression plot showing a strong correlation between phonetic subscore and cluster 1. G. Regression plot showing a perfect correlation between prosodic subscore and cluster 2.

**Figure 2.**
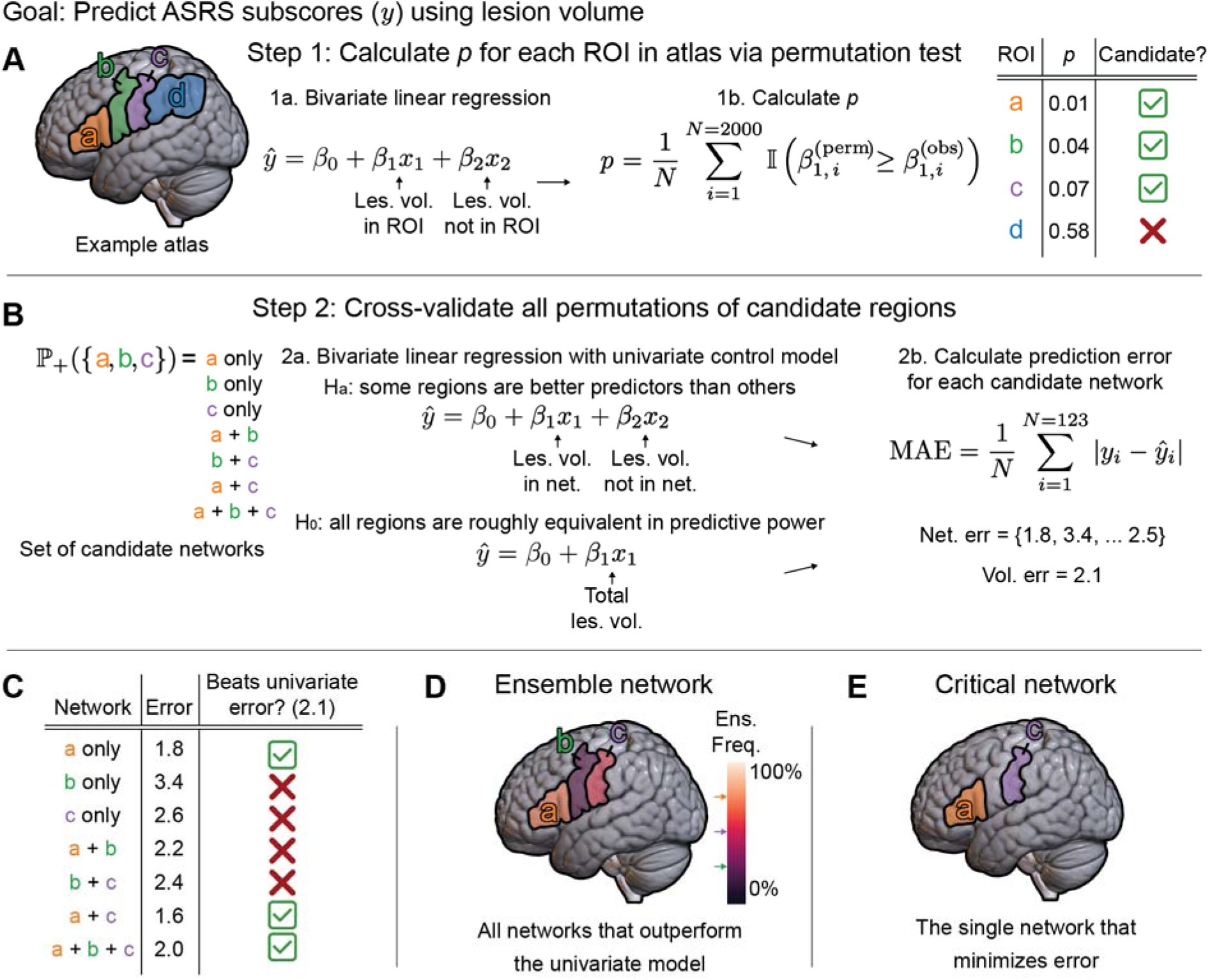
Schematic of critical network lesion-symptom mapping. Figure reprinted with permission from Walker et al., in prep. The goal of CNLSM is to predict behavioral scores from lesion data. The core assumption of the method is that some regions are more useful than others in predicting behavioral scores. A: Each atlas region is evaluated through a permutation test which selects a set of candidate regions that meet a specified *p*-value threshold. B: Candidate networks are formed from all possible combinations of candidate regions. Prediction accuracy for each candidate network is evaluated via leave-one-out cross-validation. For each network, a bivariate model is fit where behavioral scores are modeled using in-network lesion volume (first coefficient) and out-of-network lesion volume (second coefficient). C: The mean absolute error of each candidate network is tested for significance against the MAE of a univariate model predicting behavioral scores using overall lesion volume. D: The errors from each network that outperforms the univariate model are averaged together to form an ensemble model from which individual region co-occurrence and rate of prevalence within the ensemble networks can be calculated. E: The single network that minimizes error relative to the univariate and ensemble prediction errors is declared the critical network.

Because our research question concerns prosodic and phonetic abilities, we opted to use our generated ASRS-3.5-style prosodic and phonetic subscores in subsequent analyses; however, we wished to provide additional validation of the usage of these subscores in our data. To that aim, we calculated itemwise correlations between all ASRS scores and then hierarchically clustered them using an agglomerative method. Aggregating scores within these clusters allowed us to generate “subscores” without a priori assumptions about the relative subdomains of importance in AOS. We next generated a bootstrap distribution (2000 iterations) where data were shuffled prior to hierarchical clustering to estimate within-branch item co-occurrence frequency (Figure 1C). The two largest clusters closely aligned with our proposed prosodic and phonetic subscores based on the ASRS-3.5 subscores: the first cluster correlated with the phonetic subscore (*r* = 0.95; Figure 1F) and the second contained the exact same items as the prosodic subscore (*r* = 1; Figure 1G). Because our a priori categories of “prosodic subscore” and “phonetic subscore” mapped so strongly onto itemwise correlations in the data revealed through unsupervised clustering, we believe that updating ASRS-1 scores to reflect ASRS-3.5 subscores is a valid approach.

### Image Acquisition

Structural MRI for participants in the POLAR database were acquired on a Siemens 3T Prisma Fit scanner using a 20-channel head coil at the McCausland Center for Brain Imaging at the University of South Carolina. T1 and T2 images were acquired with a voxel size of 1 mm^3^. Lesions were manually demarcated on the T2 images by a licensed neurologist. T1 images and lesion masks were nonlinearly warped to the mni152 reference space to facilitate generalization across subjects via in-house scripts and SPM12. The final warped lesions and corresponding images used in lesion-symptom mapping had an array shape of 207 x 256 x 215 and a voxel size of 0.737 mm^3^.

### Lesion-symptom mapping critical networks for phonetic and prosodic ability

Brain-behavior relationships were evaluated using a novel lesion-symptom mapping technique from our group called critical network lesion-symptom mapping (CNLSM, Walker et al., in prep.). Conventional voxel-based lesion symptom mapping approaches (erroneously) assume independence between all voxels of the brain as they are statistically a series of mass univariate comparisons (Mah et al., 2014). Multivariate approaches (e.g., support vector regression) do not make such an assumption but carry their own caveats concerning interpretability and causality (Sperber, 2020). The method we employ aims to identify critical networks supporting performance on a behavioral assessment using a prediction-based framework in an attempt to mitigate these issues with common lesion-symptom mapping approaches. Lesion data are first parcellated into regions based on a functional or anatomical atlas. From the atlas, regions that are good candidates for supporting the behavior of interest (in our case, prosodic and/or phonetic ASRS subscore performance) are identified via a *p* value threshold, calculated via resampling methods. This *p* value is calculated as:

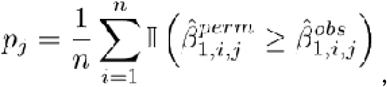

or, for a brain region *j*, the proportion of times over *n* (2000) iterations of permuted data that the β_1_ coefficient (in-ROI lesion volume) in an ordinary least squares regression of in-ROI and out-of-ROI (β_2_) lesion volume is greater than the observed β_1_ in the non-permuted data. We chose a more lenient *p* < 0.1 threshold for candidate regions to reduce the likelihood of false negatives; goodness of fit was further evaluated using techniques described below that ideally would eliminate potential false positives from use of a more lenient *p* value in this first step.

After identification of candidate regions, combinations of these regions are tested for prediction accuracy using leave-one-out cross-validation (LOOCV) on a simple multivariate linear model:

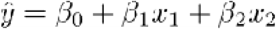

This multivariate model is identical to the one used to calculate individual brain region significance, except now β_1_ is the lesion volume within the candidate network (cf. within an individual region), and β_2_ is the lesion volume outside the candidate network (cf. outside an individual region). Mean absolute error (MAE) between predicted and actual behavioral scores was selected as the error metric and calculated as:

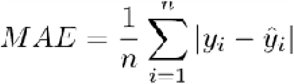

MAE for every combination of candidate regions (e.g., for 10 candidate regions, 2^10^ – 1 combinations were tested) was compared to MAE for a simple bivariate model with total lesion volume as the only predictor using a related samples *t*-test. The prose interpretation of this *t*-test is an evaluation of whether treating a set of brain regions as a distinct network within the lesioned area increases the ability of the model to predict ASRS subscores. All non-zero combinations of candidate regions were tested and any combination of candidate regions that yielded *p* < 0.05 on the *t*-test was treated as a candidate network for supporting the function. The putative “best” combination of candidate regions would be the one that minimizes MAE; that network is referred to as the “critical network.”

While a single network of brain regions best modeling the ASRS subscore of interest is identified, a subset of networks that significantly outperform the bivariate lesion volume control model while not significantly under-performing the critical network are included in analysis as the “ensemble network.” A metric of ensemble prediction across all networks in the ensemble was calculated by averaging error (i.e., equivalent to averaging the predictions and then calculating the error) so that the gains in predictive power over an ensemble of “good enough” networks of regions could be reported alongside the singular critical network. We report the critical network for each ASRS subscore modeled in our results, but we also opt to report co-occurrence of individual regions within ensemble networks and inclusion rate of individual regions within the ensemble networks to emphasize that lesion-to-symptom mappings are not monolithic. While it is often combinations of anatomically distributed regions in tandem that critically give support to a behavioral function, given the combinatoric complexity of the possible underlying networks, there are typically multiple combinations of regions that can plausibly link a set of lesion data and a set of behavioral data. Statistics derived from the ensemble network allow us to consider each region’s importance across this “multiverse” of possible explanations for our data. Individual atlas regions that have a high rate of prevalence within the ensemble networks and/or strong co-occurrence with other candidate regions in the ensemble, even if these regions are potentially absent from the singular critical network, are likely still of importance in supporting the behavioral ability. Variability in lesion-behavior mapping results across studies is quite high (Teghipco et al., 2024), which was a primary motivation for utilizing a more descriptive method of lesion-symptom mapping in CN-LSM, which identifies a set of plausible networks underlying a function in addition to the classic definitive set of regions as is presented in conventional lesion-symptom mapping methods.

Because CN-LSM is fundamentally an atlas-based method, and because different atlases can yield different findings based on their particular regional partitions, we chose to model the prosodic and phonetic ASRS subscores using two atlases and then compare results across the models. The first atlas we used was HCPex, an extension of the Human Connectome Project atlas with extended coverage of subcortical areas. HCPex is a multimodal parcellation of cortical and subcortical regions, defined through structural imaging (T1/T2 weighted thickness/myelin maps), resting state fMRI connectivity, task-based activation, and topographic organization (Huang et al., 2022). We chose HCPex because of its clear delineation of area 55b, an area of interest for apraxia of speech with anatomical proximity to the dorsal precentral speech area (Chang et al., 2020; Glasser et al., 2016; Hickok et al., 2023). Because the HCPex atlas does not contain white matter tractography, and because connectivity between the dPCSA/vPCSA and STG/aSMG (respectively) are likely important components of the prosodic and phonetic networks per prior functional imaging research (Burns et al., 2025; Hickok et al., 2023), we chose the AALCAT atlas supplied with NiiStat (https://www.nitrc.org/projects/niistat/) as our second atlas. The AALCAT atlas is a combination of the AAL cortical atlas (Collins et al., 1998) and the Catani tractography atlas (Catani & Thiebaut de Schotten, 2008).

## Results

This study aimed to identify patterns of left hemisphere post-stroke impairment associated with prosodic and phonetic processing in a cohort of people with aphasia and apraxia of speech in an effort to delineate separate neural systems associated with those functions. We employed structural MRI and a series of low-dimensional regression models to associate specific brain regions with prosodic and phonetic ability. The output of this approach is an *ensemble network* of brain regions that, when lesioned, predict prosodic and/or phonetic ability. The single optimal (in terms of prediction accuracy) network within the ensemble is deemed the *critical network.* Scores on the Apraxia of Speech Rating Scale (ASRS) were split into prosodic and phonetic subscores which served as the behavioral variables of interest (Duffy et al., 2023).

Consistent with both the PPAOS literature on prosodic and phonetic subtypes and the theoretical dual motor coordination model, we identified separable neural substrates for phonetic and prosodic ability. The patterns of frequency and co-occurrence within the ensemble networks revealed via critical network lesion-symptom mapping reflects two separable networks for prosodic and phonetic ability, with the former being more dorsal and the latter being more ventral in distribution.

The ensemble of networks supporting prosodic function localized to the frontal operculum, posterior inferior frontal junction (pIFJ; anterior to dPCSA, but within the dorsal hierarchy), dorsal somatosensory cortex (Brodmann areas 2, 3a), piriform cortex, and putamen (Table 2; Figure 3A). 40% (*n* = 13162) of the permuted networks of candidate regions predicted prosodic subscores better than the bivariate control model. In addition to the ensemble networks, the single optimal network is displayed in Table 2. While a region approximating the dPCSA was present in the ensemble networks, ventral sensorimotor regions also appear to be of importance in modeling prosodic ability. By measuring the degree of co-occurrence within ensemble networks for prosodic subscore, we saw that two regions absent from the critical network were nonetheless important in predicting prosodic ability: Brodmann area 2 (primary somatosensory cortex) and the putamen were present in 75% and 76% of ensemble networks (respectively) and frequently co-occurred with every other region found in the ensemble networks. Full maps of region frequency and co-occurrence are shown in Figures 3B and 3C, respectively.

**Figure 3.**
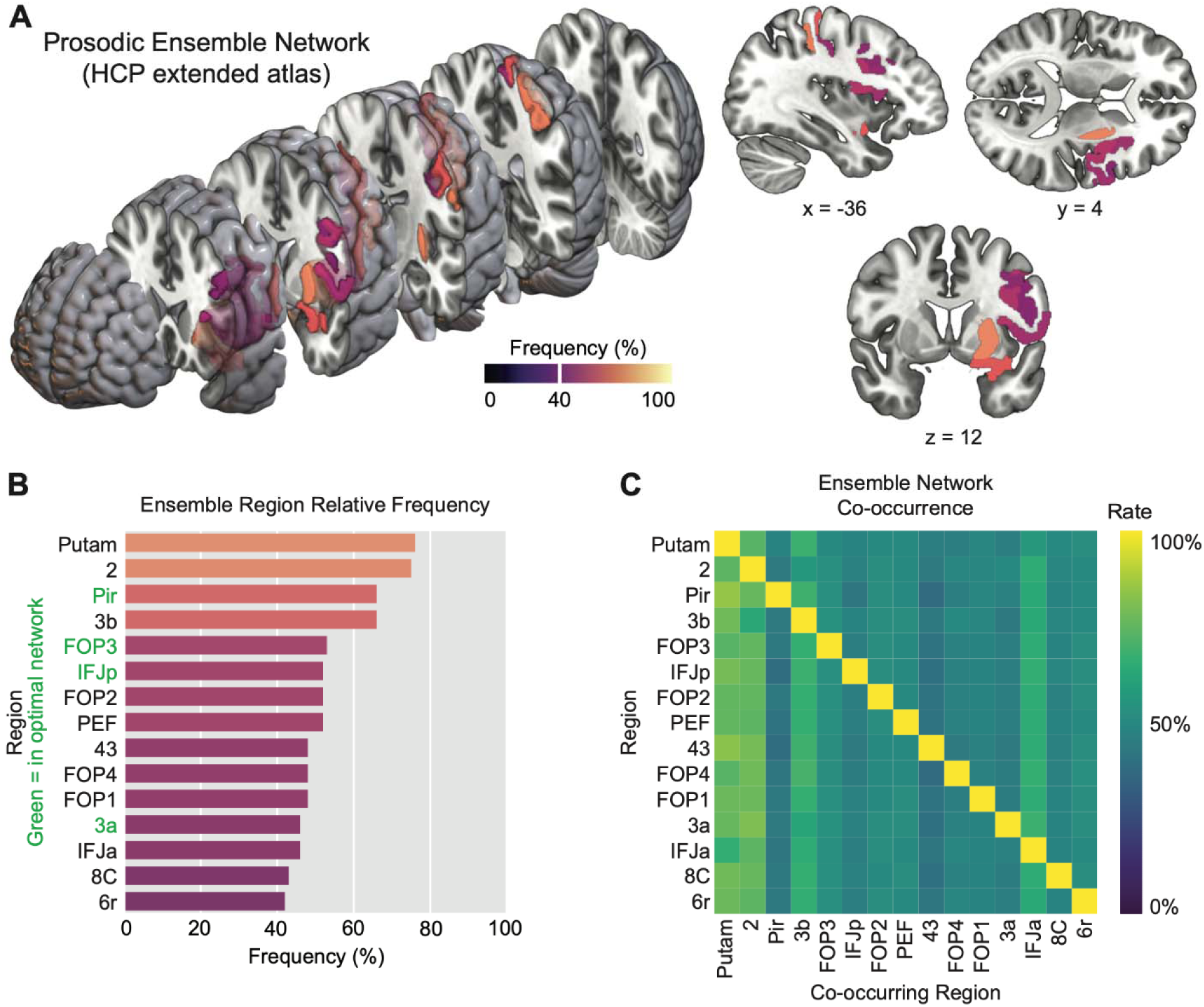
Ensemble of networks of cortical and subcortical regions supporting prosodic function. A: Left: Critical regions displayed on a 3D exploded view of an MNI template brain. Region color corresponds to relative frequency displayed in panel B. Right: Representative 2D slices and coordinates. B: Relative frequency of individual regions within the ensemble network. Color map is inherited from panel A. Region labels in green were members of the single optimal network within the greater ensemble. C: Matrix showing co-occurrence rate of individual dyads of regions within the ensemble network.

**Table 2.** Single optimal network summaries for prosodic and phonetic ASRS subscores in the HCPex atlas.

| ASRS<br>Subscore | Critical Network |  | CN<br>MAE (*p*) | Ensemble<br>MAE (*p*) | Volume MAE<br>(Univariate) |
| --- | --- | --- | --- | --- | --- |
|  | ROI | Description |  |  |  |
| Prosodic | 3a | Brodmann area 3a,S1 | 3.713<br>(0.003) | 3.847<br>(0.009) | 4.230 |
|  | FOP3 | Frontal operculum |  |  |  |
|  | IFJp | Posterior inferior frontal junction |  |  |  |
|  | Pir | Piriform cortex |  |  |  |
| Phonetic | 3a | Brodmann area 3a | 3.554<br>(*<0.001*) | 3.708<br>(*<0.001*) | 4.339 |
|  | FEF | Frontal eye fields |  |  |  |
|  | OP1 | Parietal operculum, S2 |  |  |  |
|  | OP2-3 | Parietal operculum, vestibular cortex |  |  |  |
|  | FOP2 | Frontal operculum |  |  |  |
|  | FOP3 | Frontal operculum |  |  |  |
|  | IFJp | Posterior inferior frontal junction |  |  |  |

The ensemble of networks supporting phonetic function revealed a high degree of overlap between Brodmann area 3a (primary somatosensory cortex; in 86% of ensemble networks) and every other region (Figure 4C). Relatively high ensemble network overlap was also present in the frontal/parietal opercula and also areas in granular and middle insula. 17% (*n* = 5631) of networks predicted phonetic subscores better than the control model. Despite prosodic and phonetic function localizing to separate networks of brain regions, we observed a moderate amount of overlap in the frontal operculum, pIFJ, and primary somatosensory cortex. The single optimal network is displayed in Table 2.

**Figure 4.**
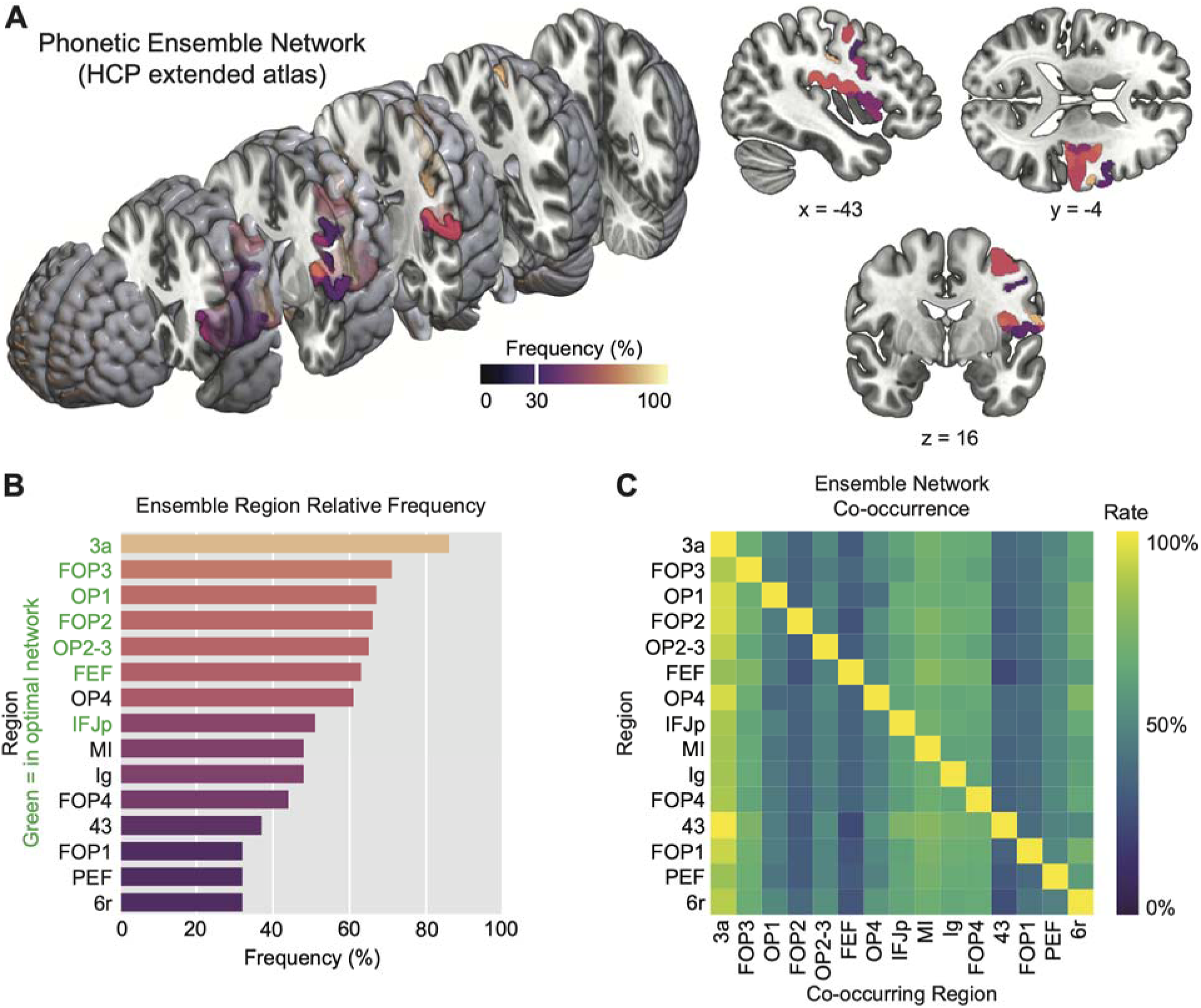
Ensemble of networks of cortical and subcortical regions supporting phonetic function. A: Left: Critical regions displayed on a 3D exploded view of an MNI template brain. Region color corresponds to relative frequency displayed in panel B. Right: Representative 2D slices and coordinates. B: Relative frequency of individual regions within the ensemble network. Color map is inherited from panel A. Region labels in green were members of the single optimal network within the greater ensemble. C: Matrix showing co-occurrence rate of individual dyads of regions within the ensemble network.

Another set of networks were identified using the AALCAT atlas (Catani & Thiebaut de Schotten, 2008; Collins et al., 1998), which contains both cortical and white matter regions of interest, to better describe the medial regions of interest identified using the HCPex atlas (Table 3; Figure 5). Ensemble network results in the AALCAT atlas reflect the expected connectivity of prosodic (auditory cortex, laryngeal motor control) and phonetic (supramarginal gyrus, Spt) ability (Figure 5A). For prosodic subscore, the ensemble network consisted of 61.7% (*n* = 2529) of permuted networks and the ensemble network for phonetic subscore consisted of 38.9% (*n* = 397) of permuted networks. For prosodic function, the corticospinal tract (68% of ensemble networks) frequently co-occurred with almost all other candidate regions (Figure 5C). Insular white matter tracts were also relatively common in the prosodic ensemble networks (62% of ensemble networks) and disproportionately co-occurred with precentral tracts compared to other candidate regions. Overall, prosodic subscore localized to a large network connecting the frontal/central opercula, postcentral gyrus, insula, cerebellum, and temporal lobe. The patterns of co-occurrence and frequency in the prosodic ensemble network suggests that prosodic function is supported by connections spanning between dorsal regions in somatosensory and motor cortex and auditory regions supporting auditory feedback control, but also connections between dorsal sensory/motor cortex and the cerebellum/corticospinal tract supporting primary efferent control of the larynx.

**Figure 5.**
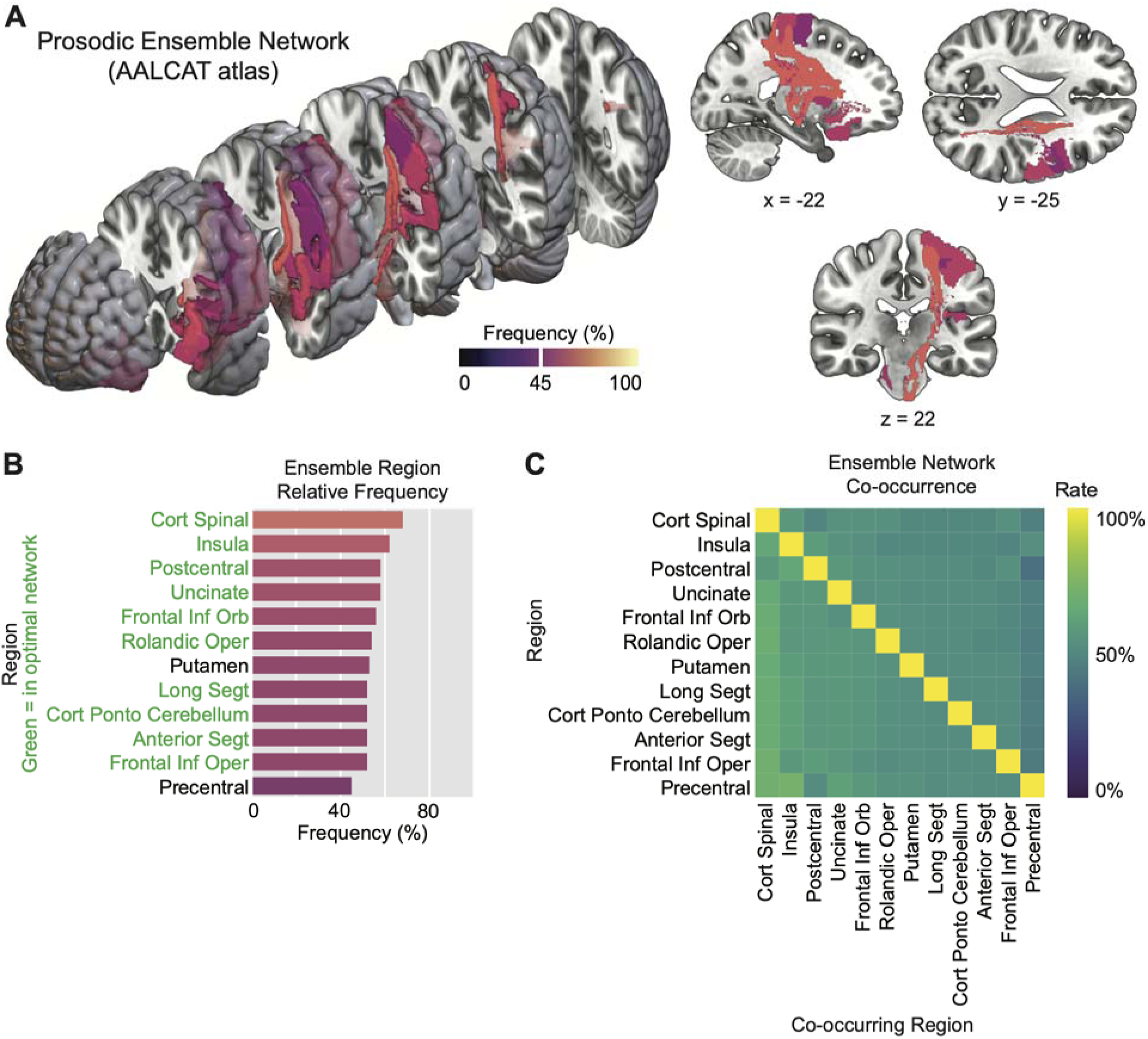
Ensemble of networks in an atlas containing white matter tracts supporting prosodic function. A: Left: Critical tracts displayed on a 3D exploded view of an MNI template brain. Region color corresponds to relative frequency displayed in panel B. Right: Representative 2D slices and coordinates. B: Relative frequency of individual tracts within the ensemble network. Color map is inherited from panel A. Tracts in green were members of the single optimal network within the greater ensemble. C: Matrix showing co-occurrence rate of individual dyads of tracts within the ensemble network.

**Table 3.** Single optimal network summaries for prosodic and phonetic ASRS subscores in the HCPex atlas.

| ASRS Subscore | Critical Network | CN MAE (*p*) | Ensemble MAE (*p*) | Volume MAE (Univariate) |
| --- | --- | --- | --- | --- |
| Prosodic | Frontal Inf Oper | 3.778 (0.005) | 3.845 (0.005) | 4.230 |
|  | Frontal Inf Orb |  |  |  |
|  | Rolandic Oper |  |  |  |
|  | Insula |  |  |  |
|  | Postcentral |  |  |  |
|  | Anterior Segment |  |  |  |
|  | Cortico Ponto Cerebellum |  |  |  |
|  | Cortico Spinal |  |  |  |
|  | Long Segment |  |  |  |
|  | Uncinate |  |  |  |
| Phonetic | Anterior Segment | 3.547 (<0.001) | 3.707 (<0.001) | 4.339 |
|  | Long Segment |  |  |  |

Phonetic subscore, on the other hand, localized to a smaller network consisting primarily of the arcuate fasciculus, which runs between ventral sensorimotor cortex (vPCSA) and area Spt (Figure 6). Co-occurrence with other regions was also strongest for the arcuate fasciculus (87% of ensemble networks), which is in line with the regions identified in the single optimal network (Table 3) and emphasizes the importance of connectivity between ventral motor control regions and auditory-to-motor transformation regions in the inferior parietal lobe and temporoparietal junction. While we did not see a smaller, more focal network supporting prosody as observed in neurodegenerative work (Josephs et al., 2013; Utianski et al., 2018), the critical networks identified for phonetic and prosodic subscores appear to support the theory that phonetic ability in vPCSA is functionally connected to the anterior supramarginal gyrus while prosodic ability in dPCSA is connected to auditory cortex (Burns et al., 2025; Hickok et al., 2023).

**Figure 6.**
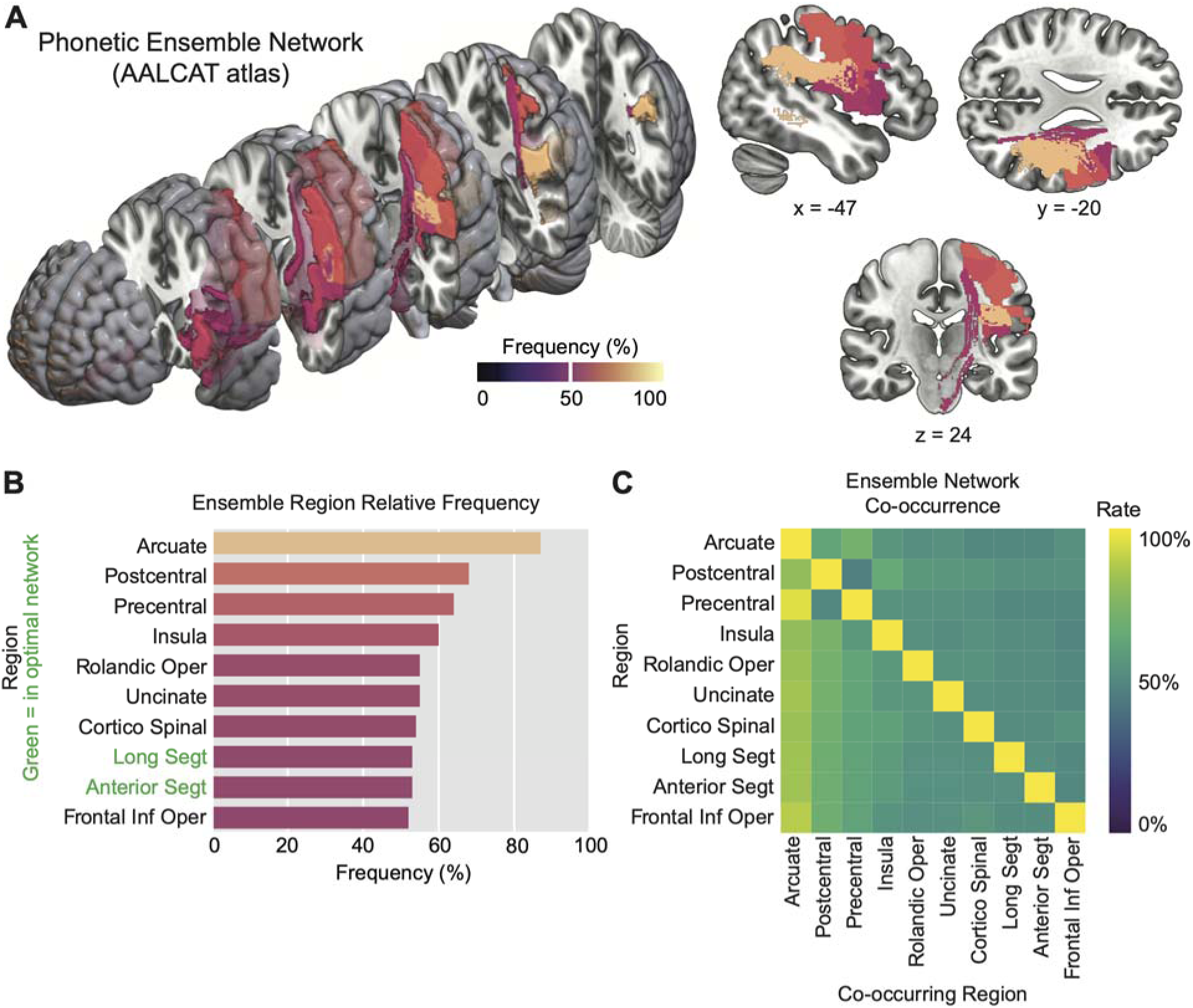
Ensemble of networks in an atlas containing white matter tracts supporting phonetic function. A: Left: Critical tracts displayed on a 3D exploded view of an MNI template brain. Region color corresponds to relative frequency displayed in panel B. Right: Representative 2D slices and coordinates. B: Relative frequency of individual tracts within the ensemble network. Color map is inherited from panel A. Tracts in green were members of the single optimal network within the greater ensemble. C: Matrix showing co-occurrence rate of individual dyads of tracts within the ensemble network.

## Discussion

This study demonstrates that the prosodic and phonetic components of apraxia of speech, as quantified by subscores on the Apraxia of Speech Rating Scale, localize to separate neural pathways in a large cohort of left-hemisphere stroke survivors with aphasia and/or apraxia of speech. We utilized a probabilistic lesion-symptom mapping technique that aimed to identify the relative likelihood that individual brain regions are important in modeling prosodic and/or phonetic function. This technique generated an ensemble of brain networks that, when treated as functionally distinct from overall lesion volume, plausibly improved the ability to predict behavioral scores from lesion data. Our use (and prior development) of a prediction-based, ensemble approach to lesion-symptom mapping was motivated by the inherent noise of lesion studies. While conventional lesion-symptom mapping techniques identify a single solution for mapping brain to behavior, the prediction-based nature of our mapping technique yields insights into other maps that still plausibly model the brain-behavior relationship despite not being a single most-optimal mapping of brain-to-behavior.

The ensemble of brain regions that best predicted the prosodic subscore of the ASRS consisted of central sensory and motor areas, subcortical motor nuclei, the cerebellum and cortico-cerebellar pathways, and white matter connections between primary sensorimotor cortex and auditory cortex/insula. The ensemble of brain regions that best predicted the phonetic subscore of the ASRS consisted of partially overlapping central sensory and motor areas alongside the dorsal aspect of the arcuate fasciculus connecting ventral sensorimotor and inferior parietal cortex. These results support the interpretation that prosodic and phonetic deficits in speech output have distinct neural foundations and implies that considering prosodic/phonetic impairments as discrete treatment goals during rehabilitation may improve patient outcomes in motor coordination disorders such as apraxia of speech.

While we initially hypothesized that prosodic ability would localize to the lateral cortical surface in the dorsal precentral speech area (dPCSA) and phonetic ability would localize to a more distributed frontoparietal network including the ventral PCSA, the networks identified by our approach paint a more complex picture. Rather than mapping to discrete lateral cortical areas, prosodic and phonetic ability as determined by ASRS subscores mapped instead onto an overlapping mixture of cortical and subcortical areas in the HCPex atlas. Phonetic subscore localization partially overlapped with prosodic subscore in the frontal operculum, primary sensory cortex, and posterior IFG. While we interpret these networks as partially overlapping yet still distinct, in the context of our results both prosodic and phonetic function could be described as dorsal *and* ventral. We identified a myriad of regions outside of the precentral gyrus that played an important role in modeling prosodic and phonetic function. Brodmann area 2, part of primary somatosensory cortex, was present in the ensemble networks for prosodic and phonetic function. For prosody, BA2 likely plays a role in proprioceptive monitoring of pitch, a necessary component of feedback control during speech (Houde & Nagarajan, 2011), while its role in phonetic function is likely related to somatosensory feedback during articulation. Future experimental work using high-temporal-resolution electrophysiological recordings could confirm a post-articulatory role for BA2 in prosodic and phonetic function. The putamen and piriform cortex also emerged as important regions supporting prosodic function. The putamen is a subcortical motor control nucleus involved in pitch regulation during singing (Zarate & Zatorre, 2008), while piriform cortex is less directly related to prosodic function in the literature. It is possible the involvement of piriform cortex is due to its medial proximity to auditory regions in the insula and planum temporale, meaning the inclusion of piriform cortex in the optimal network for prosody may be reflective of connectivity to auditory regions supporting prosodic function. Lastly, insular regions were present in both prosodic and phonetic ensemble networks, but absent from the optimal networks for both ASRS subscores. The superior precentral gyrus of the insula in particular has been linked to apraxia of speech, specifically in an articulatory role, in prior literature, which supports the insular mapping of phonetic function in the current study ((Baldo et al., 2011; Dronkers, 1996); but see also disagreement in (Fedorenko et al., 2015)).

There is no direct link between prosody and the insula in the literature, but limited studies have shown a role for the insula in auditory processing during speech production (Kurteff et al., 2024; Woolnough et al., 2019). The insula is a multifunctional brain region that plays a role in numerous sensory, motor, and cognitive processes (Kurth et al., 2010) and the vast majority of middle cerebral artery strokes implicate the insula in some form (Hillis et al., 2004) which makes further interpretation of these results (an atlas-based lesion study) particularly difficult. In general, differences between underlying patterns of impairment in stroke and in neurodegeneration may explain differences between the current study and prior work done in progressive aphasias and AOS.

The white matter tracts in our mapping of the AALCAT atlas more clearly separated prosodic and phonetic processing into distinct pathways. We interpret the white matter pathways supporting prosodic function as reflecting connectivity of primary sensorimotor cortex to auditory regions (as proposed by Hickok et al., 2023) as well as subcortical and cerebellar motor control regions. A large white matter network supporting prosody may reflect the increased demands on sensory systems supporting prosody compared to articulation, as vocal feedback monitoring is an important component of prosodic control. The involvement of more long-distance fibers such as the corticopontocerebellar and corticospinal tracts supports this hypothesis. The corticospinal tract involvement also likely reflects the role primary efferent control of the larynx plays in prosodic control and the anatomical proximity of dorsal laryngeal motor cortex (Dichter et al., 2018) and the dPCSA. The most important white matter pathways for modeling phonetic function were the anterior and long segments of the arcuate fasciculus, which connects the inferior frontal gyrus and ventral precentral gyrus to auditory-motor regions in the Sylvian-parietal-temporal (Spt) junction. This reinforces the connections proposed in the Hickok et al., 2023 dual motor coordination: an auditory-to-precentral pathway supports prosodic function and a parietal-to-precentral pathway supports phonetic function.

There are several plausible explanations for why the results of the current study are not a clear-cut validation of the dPCSA/vPCSA dichotomy put forth in Hickok et al., 2023. Firstly, we observed some collinearity in our prosodic and phonetic subscores (Figure 1D, *r* = 0.52). This does not imply that prosodic and phonetic function cannot be separated using a scale such as the ASRS, but rather highlights that at a certain degree of severity, patients with expressive speech deficits will score poorly on *both* prosodic and phonetic subcomponents of the ASRS. For mild-to-moderate cases, there are individual subjects who clearly are selectively impaired in a single domain, suggesting that prosodic and phonetic ability are separable using the ASRS. It is possible that the partial overlap we identified in networks supporting prosodic and phonetic function is related to this collinearity in more severe cases but we lack sufficient power to do a follow-up analysis in subsets of our cohort. Therefore, it is difficult to conclude in the present study whether the partial overlap in prosodic and phonetic networks is due to shared neural substrates or an artifact of more severe cases in the cohort. Similarly, we could not construct ensemble network statistics for the subset of our cohort that had a diagnosis of AOS (*n* = 61) for power reasons. Our results could also highlight a fundamental limitation of lesion-symptom mapping studies in that these are noisy data, especially in comparison to our benchmark of Hickok et al., 2023, which is a literature review. We anticipate future studies of prosodic and phonetic function will provide converging evidence that support the separation of these processes into separate anatomical hubs within precentral cortex and their corresponding underlying pathways.

In terms of translating these results to the clinic, more work is of course needed. However, we advocate that clinicians assessing and treating post-stroke AOS pay attention to whether patients make errors that fall primarily into a prosodic or phonetic pathology, as the results presented in this study support at least a partial neurobiological separation of these faculties. Future studies should aim to better categorize prosodic and phonetic deficits in post-stroke AOS, but behavioral guidelines from the PPAOS literature offer a good starting point (Utianski et al., 2018): prosodic post-stroke AOS is likely marked by difficulty with syllable segmentation and reduction of words per breath group, while phonetic post-stroke AOS is likely marked by speech sound substitutions and distortions.

## Conclusion

This study provides motivation for conceptualizing post-stroke apraxia of speech as two separate profiles of impairment with different neural foundations in a fashion that parallels such a split in the primary progressive AOS literature. *Prosodic* AOS is neurobiologically marked by impairment in a precentral-to-auditory motor coordination stream while *phonetic* AOS is marked by impairment in a precentral-to-parietal motor coordination stream. The existence of separable neural pathways for these aspects of speech production in a cohort of stroke survivors with aphasia and/or AOS supports the theoretical separation of motor coordination into two hierarchies governing laryngeal (or prosodic) and supralaryngeal (or phonetic) coordination.

## Acknowledgements

We thank the participants from the POLAR study at C-STAR for their generous participation in this research. This study was supported by a P50 grant from the NIDCD (DC014664; PI Julius Fridriksson).

## Data Availability Statement

In accordance with the National Institutes of Health policy for data sharing (https://grants.nih.gov/policy-and-compliance/nihgps), upon completion of the POLAR trial and dissemination of primary study results, these data will be made available to the public. The data that support the findings of this study and the code to generate the results are available from the corresponding author upon request.

